# Multi-Method Molecular Characterisation of Human Dust-Mite-associated Allergic Asthma

**DOI:** 10.1101/446427

**Authors:** E. Whittle, M.O. Leonard, T.W. Gant, D.P Tonge

**Affiliations:** School of Life Sciences, Faculty of Natural Sciences, Keele University, ST5 5BG; Centre for Radiation, Chemical and Environmental Hazards, Public Health England, OX11 0RQ

## Abstract

Asthma is a chronic inflammatory disorder of the airways. Disease presentation varies greatly in terms of cause, development, severity, and response to medication, and thus the condition has been subdivided into a number of asthma phenotypes. There is still an unmet need for the identification of phenotype-specific markers and accompanying molecular tools that facilitate the classification of asthma phenotype. To this end, we utilised a range of molecular tools to characterise a well-defined group of adults with poorly controlled asthma associated with house dust mite (HDM) allergy, relative to non-asthmatic control subjects. Circulating messenger RNA (mRNA) and microRNA (miRNA) were sequenced and quantified, and a differential expression analysis of the two RNA populations performed to determine how gene expression and regulation varied in the disease state. Further, a number of circulating proteins *(IL-4, 5, 10, 17A, Eotaxin, GM-CSF, IFNy, MCP-1, TARC, TNFa, Total IgE, and Endotoxin)* were quantified to determine whether the protein profiles differed significantly dependent on disease state. Finally, assessment of the circulating “blood microbiome” was performed using 16S rRNA amplification and sequencing. Asthmatic subjects displayed a range of significant alterations to circulating gene expression and regulation, relative to healthy control subjects, that may influence systemic immune activity. Notably, several circulating mRNAs were detected in the plasma in a condition-specific manner, and many more were found to be expressed at altered levels. Proteomic analysis revealed increased levels of inflammatory proteins within the serum, and decreased levels of the bacterial endotoxin protein in the asthma state. Comparison of blood microbiome composition revealed a significant increase in the Firmicutes phylum with asthma that was associated with a concomitant reduction in the Proteobacteria phylum. This study provides a valuable insight into the systemic changes evident in the HDM-associated asthma, identifies a range of molecules that are present in the circulation in a condition-specific manner (with clear biomarker potential), and highlights a range of hypotheses for further study.

## Introduction

Asthma is a chronic inflammatory disorder of the airways and is a global public health concern due to increasing prevalence and mortality rates (1–4). The World Health Organisation has estimated that 300 million people are living with asthma, and that 250,000 individuals die prematurely each year as a result of the disease (5).

Asthma can develop during childhood (early-onset) or in adulthood (late-onset) and is characterised by chronic inflammation of the airways and intermittent episodes of reversible airway obstruction (6,7). Over time, chronic inflammation of the airways results in airway hyper-responsiveness and structural changes, including airway fibrosis, goblet cell hyperplasia, increased smooth muscle mass, and increased angiogenesis (7,8).

The causes of asthma are multifactorial, and include a complex variety of environmental, immunological, and host genetic factors (7,9–13). Disease typically occurs in genetically predisposed individuals (13,14), and clinical presentation is highly heterogenous (15). Disease can vary greatly in terms of disease onset and response to treatment (16). It can present as a chronic, stable disease, but also as intermittent asthma exacerbations that can be fatal (17). Symptoms can be mild or severe and arise as a result of a multitude of factors, including immunoglobulin-E (IgE) mediated allergic responses, exposure to pollutants, exercise, stress, or airway infections (17).

The complex nature of asthma pathogenesis has resulted in speculation as to whether asthma is a single disease, or a spectrum of related diseases with subtle but distinct differences in aetiology and pathophysiology (18,19). This has led to asthma being separated into a number of phenotypes, which are then further subdivided into several endotypes (6,15,18–20). These asthma phenotypes are triggered by complex gene-environment interactions and respond differently to the various asthma medications available. Individuals with eosinophilic asthma, for instance, have been reported to have a good therapeutic response to inhaled or oral corticosteroid therapy, whereas individuals with neutrophilic asthma have been found to respond poorly to this therapeutic approach (21).

Diagnostic tools for identifying the various asthma phenotypes are limited, and thus optimal treatment protocols are not being utilised in a number of patients. Moreover, despite decades of research, there has been little progress in the development of treatments since the introduction of inhaled ß2 adrenoceptor 2 selective agonists (1969) and inhaled glucocorticosteroids (1974) (15). Long-term use of these medications has been associated with a number of health concerns (22), including the stunting of growth in children (23), cataract development (24,25), osteoporosis (26,27), and cardiovascular events (28). Overall, an estimated 5-10% of asthmatics fail to respond to conventional medications (29). In order to improve patient response to treatment, and / or assist in the development of new therapeutics, an improved knowledge of the molecular mechanisms that underlie the various asthma phenotypes is required. Long-term, this may also facilitate the targeted use of conventional asthma therapies, and facilitate the development of new medications aligned to the individual asthmatic phenotypes, subsequently reducing asthma mortality and improving quality of life.

The focus of this study was to characterise, at the molecular level, a small but well-defined cohort of patients with atopic asthma associated with house dust mite (HDM) allergy. Global estimates suggest that 1-2% of the world’s population are sensitive to HDM (30), as are approximately 50% of asthmatic patients (30,31). HDM sensitivity has been linked to increased asthma severity (32) and almost one-third of patients with HDM sensitivity are unresponsive to current asthma therapies (33). Increasing our understanding of this specific asthma phenotype is therefore crucial. To this end we performed a comprehensive molecular characterisation of (1) circulating mRNAs, (2) circulating microRNAs, (3) circulating protein-based markers of the immune response and (4) integrated these data with our previous work characterising evidence of a circulating microbiome.

## Methods

### Donor Population

Atopic asthmatic individuals (n=5) with physician-diagnosed HDM allergy, and gender and age-matched healthy control subjects (n=5) were recruited to the study via SeraLabs Limited. Asthma patients were selected on the basis that they had developed atopic asthma during early childhood and that their condition had continued into adulthood and remained “poorly controlled”. A full list of recruitment criteria is presented in **(Table 1).**

**Table 1:** Donor population characteristics required for the study

| <u>**Patient Criteria**</u> |
| --- |
| • Have a BMI < 30<br>• Be a non-smoker<br>• Have been diagnosed with atopic asthma during childhood<br>• Have severe/ poorly controlled asthma<br>• Must not be on any oral steroid treatment<br>• Must be allergic to the house dust mite<br>• Must not have diabetes, COPD, or hypertension |

Whole blood was drawn, following alcohol cleansing of the skin surface, into EDTA containing tubes and stored on ice prior to centrifugation at 1000×g to obtain the plasma component. All samples were analysed anonymously, and the authors obtained ethical approval and written informed consent to utilise the samples for the research reported herein.

The Independent Investigational Review Board Inc. ethically approved sample collection by Sera Laboratories Limited from human donors giving informed written consent. Furthermore, the authors obtained ethical approval from Keele University Ethical Review Panel 3 for the study reported herein. All experiments were performed in accordance with relevant guidelines and regulations.

### Analysis of Inflammatory proteins

Plasma levels of interleukin (IL)-4, IL-5, IL-10, IL-13, IL-17A, IFNy, TARC, Eotaxin, GM-CSF, MCP-1, RANTES, and TNFα, was determined using a qualitative enzyme-linked immunosorbent assay (ELISA) custom designed for this study. Two multianalyte sandwich ELISAs (Qiagen) were used, and analysis of the inflammatory proteins was achieved using the recommended Multi-Analyte ELISArray kit protocol (QIAGEN). Statistical analysis was performed by carrying out a Shapiro-Wilk normality test and a Wilcox rank sum test using *R* software Version 3.5.0.

### Quantitative analysis of total IgE

The concentration of total immunoglobulin E (IgE) was determined using sandwich ELISA (Genesis Diagnostics Ltd). The ELISA was performed in duplicate using the recommended protocol, and absorbance was measured at 450nm using an ELX800 absorbance reader (BioTek). Statistical analysis was performed by carrying out a Shapiro-Wilk normality test and an unpaired T test using *R* software Version 3.5.0.

### Quantitative analysis of endotoxin concentration

Circulating bacterial endotoxin concentration was measured using a Pierce™ Limulus Amebocyte Lysate (LAL) Chromogenic Endotoxin quantitative kit (Thermo Scientific). The assay was performed in triplicate using the recommended protocol, and absorbance was measured at 450nm using an ELX800 Absorbance reader (BioTek). Statistical analysis was performed by carrying out a Shapiro-Wilk normality test and an unpaired T test using *R* software Version 3.5.0.

### Total RNA extraction

Total RNA was extracted from 500μl of human plasma using the Qiagen serum and plasma miRNeasy kit. The quantity and quality of the RNA extracts was determined using the QuBit fluorimeter (Invitrogen) and BioAnalyzer (Agilent).

### Library Preparation and Next Generation Sequencing

Messenger RNA (mRNA) sequencing libraries were prepared using the SMARTer Universal Low Input RNA kit, and sequenced (Illumina HiSeq 2000) with a paired-end 90 nucleotide read metric. Small RNA sequencing libraries were prepared using the TruSeq small RNA library kit (Illumina), and sequencing was conducted on the Illumina HiSeq 2000 platform.

Raw sequencing data were trimmed of sequencing adaptors and low-quality reads removed using the Trim Galore package - a wrapper that incorporates CutAdapt and FastQC. For whole transcriptome analysis, quality-controlled reads were aligned to the Human Genome build hg19 using TopHat, a splice-junction aware mapping utility necessary for the successful mapping of intron-spanning (multi-exon) transcripts. Transcriptome assembly was performed using CuffLinks and a merged transcript representation of all samples produced using CuffMerge. Transcripts expressed at significantly different levels between the asthma and control samples were identified using CuffDiff, with a Q value ≤ 0.05 considered significant (34). MicroRNA (miRNA) analysis was performed by mapping miRNA reads to miRbase Version 21 using sRNAtoolbox (35). Differential expression of the miRNA reads was determined following statistical analysis with edgeR for R (36).

### Biological Pathway Analysis

Biological functions of the mRNA and miRNA that were differentially expressed between asthma and control subjects (defined as Q ≤ 0.05 in the mRNA dataset; and FDR ≤ 0.05 in the miRNA dataset) were determined using Ingenuity Pathway analysis (IPA) software.

Networks of genes comprising known biological processes were identified using IPA. Causal inference analysis was then applied to determine upstream regulators that may explain the pattern of differential expression seen. Casual inference analysis involved the generation of an enrichment score (Fisher’s exact test *P* value) and a Z score to determine the possible upstream biological causes of the differential gene expression observed in the asthmatic subjects (37). The enrichment score measured the overlap of observed and predicted regulated gene sets, whilst the Z score assessed the match of observed and predicted up/ down regulation patterns (37). Putative regulators that scored an overlap P value ≤ 0.05 were deemed statistically significant, and the Z scores were used to determine the activity of the putative regulators (an upstream regulator with a Z score greater than 2.0 was considered activated, whilst an upstream regulator with a Z score less than −2.0 was considered deactivated). Causal inference analysis was also used to predict the downstream effects the differentially expressed genes and miRNA could have on biological processes and functions in the asthmatic subjects.

### Circulating microbiome analysis

We have previously reported evidence of a circulating microbiome in the blood of both asthmatic and healthy patients (38) using oligonucleotide primers reported in (Supplementary Materials, S1). Here, we re-analysed this data with the aim of identifying organisms that were differentially present or abundant dependent on disease status. The QIIME pipeline was used for quality filtering of DNA sequences, demultiplexing, and taxonomic assignment. Alpha diversity was determined by calculating Shannon and Chao1 diversity indices. Differences in relative abundance was calculated by performing Shapiro-Wilk normality tests and the appropriate statistical test (unpaired T tests when the samples displayed gaussian distribution and Wilcox rank sum test when the samples did not display Gaussian distribution) on bacterial abundance data (read counts normalised to the total number of bacterial reads per patient) using *R* software Version 3.5.0.

In addition to standard statistical tests, the linear discriminant analysis effect size (LefSe) method was used to identify the bacterial taxa most likely to explain the differences in microbial populations present in the asthmatic cohort compared to the control cohort. In brief, the non-parametric factorial Kruskal-Wallis sum-rank test was applied to the 16S relative abundance data in order to detect features with significant differential abundance in the asthmatic cohort compared to the control group. A set of pairwise tests among subclasses using the unpaired Wilcoxon rank-sum test were then carried out to assess whether the detected differences in relative abundance were consistent with respect to biological behaviour. Linear discriminant analysis (LDA) was then performed to predict the effect of each identified differentially abundant bacterial taxa.

## Results

### Patient Recruitment and Characterisation

Five female asthmatic subjects were recruited in accordance with the inclusion criteria detailed in (**Table 1**). The mean age of the asthmatic subjects was 39.6 ± 11.7 years, and all had been clinically diagnosed with atopic asthma during early childhood (mean age of onset = 6.2 ± 3.2 years) (**Table 2**). At the time of sample collection, the asthmatic subjects were on prophylactic therapy to minimise the occurrence of disease symptoms (see Supplemental Material, S2). Asthma severity was determined using the internationally recognised Asthma Control Questionnaire (ACQ) (39,40), and all the asthmatic subjects scored a total > 10.0 (mean total score = 10.8 ± 0.75) (see Supplemental Material, S2). Additionally, three of the asthmatic subjects were clinically diagnosed with other atopic diseases, including allergic rhinitis, allergic dermatitis, and nasal polyps (see Supplemental Material, S2).

**Table 2:** Characterisation of the asthmatic (n = 5) and control subjects (n = 5) at the time of sample collection. S.D. = standard deviation

| Characteristic | Allergic Asthmatics | Non-Asthmatics |
| --- | --- | --- |
| **Demographic characteristics** |  |  |
| **Age - yr** |  |  |
| Mean (S.D) | 39.6 (11.7) | 39.4 (10.3) |
| Range | 19 - 52 | 23 - 49 |
| **Race or ethnic group – no. (%)** |  |  |
| Caucasian | 2 (40) | 2 (40) |
| Hispanic | 3 (60) | 3 (60) |
| **Sex – no. (%)** |  |  |
| Female | 5 (100) | 5 (100) |
| Male | 0 (0) | 0 (0) |
| **Smoking History** |  |  |
| **Smoking Status – no (%)** |  |  |
| Never Smoked | 5 (100) | 5 (100) |
| Former Smoker | 0 (0) | 0 (0) |
| Smoker | 0 (0) | 0 (0) |
| **BMI** |  |  |
| Mean (S.D) | 24.4 (2.6) | 24.3 (2.1) |
| Range | 21.5 – 27.8 | 21 – 26.4 |

Five non-asthmatic females with a mean BMI of 24.3 ± 2.1 were recruited to the study as healthy controls. The control subjects had never smoked and had a mean age of 39.4 ±10.3 years (**Table 2**). Two of the controls, Control_2 and Control_3, reported self-diagnosed dermatitis, although neither had received diagnosis by a physician for this condition.

### Inflammatory proteins

To determine the immune status of the asthmatic patients at the time of sample collection, characterisation of various chemokines and cytokines associated with asthma pathology was performed.

Qualitative ELISA was performed on the blood samples in order to profile the inflammatory state of the asthmatic and control, and inflammatory proteins under investigation included interleukin (IL)-4, IL-5, IL-10, IL-13, IL-17A, eotaxin, granulocyte-macrophage colony-stimulating factor (GM-CSF), interferon gamma (IFNy), monocyte chemoattractant protein 1 (MCP-1), thymus and activation regulated chemokine (TARC), and tumour necrosis factor alpha (TNFA). Additionally, the concentration of the pro-inflammatory bacterial endotoxin protein was measured, and total IgE present in the blood was quantified to determine the atopic state of the asthmatic subjects.

With regards to the host-derived inflammatory proteins, 10 out of the 12 inflammatory proteins under investigation were detected in the blood samples (see Supplementary Materials, S3).

Overall the asthmatic subjects were found to have elevated levels of inflammatory proteins compared to the controls, as determined by increased levels of all inflammatory proteins examined. This was particularly apparent for chemokines TARC (Fold change = 4.173; P value = 0.095), GM-CSF (Fold change = 3.607; P value = 0.111), and IFN/(Fold change = 20.871; P value = 0.195) (**Figure 1A, B, and C**). However, it should be noted that there were no statistically significant increases detected for any of the individual proteins. This was likely due to the asthmatic subjects having a greater level of diversity with regards to inflammatory protein levels compared to the control subjects (**Figure 1**).

**Figure 1:**
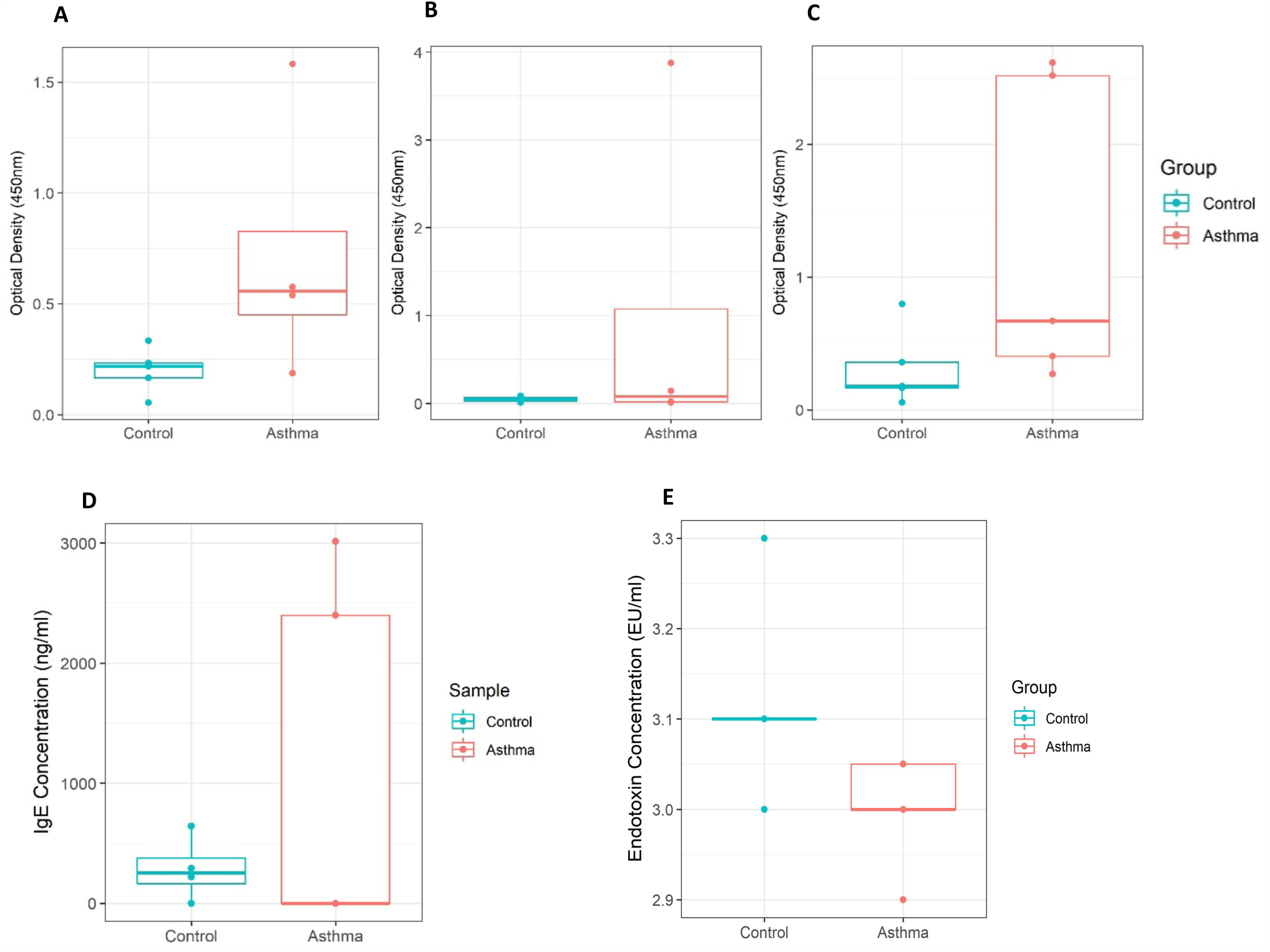
Analysis of circulatory inflammatory proteins present in blood samples from control subjects (n = 5) and asthma subjects (n = 5). **A =** levels of GM-CSF present in the blood of asthmatic subjects (n = 5) and control subjects (n = 5) using qualitative ELISA analysis, P value = 0.111 (Wilcoxon rank sum test with continuity correction); **B =** levels of IFN/present in the blood of asthmatic subjects (n = 5) and control subjects (n = 5) using qualitative ELISA analysis, P value = 0.195 (Wilcoxon rank sum test with continuity correction); **C =** levels of TARC in the blood using of asthmatic subjects (n = 5) and control subjects (n = 5) qualitative ELISA analysis, P value = 0.095 (Wilcoxon rank sum test with continuity correction); **D =** Concentrations of total IgE protein present in the blood of asthmatic subjects (n = 4) and control subjects (n = 5) using quantitative ELISA analysis, P value =1.0 (Wilcoxon rank sum test with continuity correction); **E =** Concentrations of bacterial endotoxin present in the blood of asthmatic subjects (n = 5) and control subjects (n = 5) using Limulus Amebocyte Lysate (LAL) Chromogenic quantification. P value = 0. 0650 (unpaired T test). EU/ml = endotoxin units per millilitre. Data points at 3.1 EU/ml for control = 3; Data points at 3.0 EU/ml for asthma = 2; Data points at 3.05 EU/ml for asthma = 2.

Of interest was the levels of IL-17A observed. This protein whilst not significantly increased in the asthmatic subjects (P value = 0.413), was found to be present at higher levels in asthmatic subjects who suffered additional atopic complications (Asthma_1, Asthma_2, and Asthma_4) and the two control subjects who had self-reported atopic dermatitis (Control_2 and Control_3) (see Supplementary Material S2 and S3). This suggests that whilst systemic levels of this cytokine are not elevated in asthma, IL-17A levels may be elevated in the blood of individuals with other atopic conditions, such as allergic rhinitis and atopic dermatitis.

Moreover, the asthmatic cohort appeared divided with regards to the inflammatory protein profiles, whereby asthmatic subjects Asthma_2 and Asthma_4 typically had high levels of circulatory inflammatory proteins, whilst asthmatic subjects Asthma_1, Asthma_3 and Asthma_5 displayed protein levels similar to those observed for the control subjects. This is reflective of the heterogenous nature of asthma pathology and suggests that possibility of asthma sub-phenotypes that display varying levels of circulatory inflammatory proteins. We comment upon this heterogeneity, and the impact of this upon sample size selection in the concluding section.

Total IgE was detected in 50% of the blood samples under investigation (three control subjects and two asthmatic subjects (**Figure 1D**).

For the purpose of statistical analysis, samples with undetectable levels of IgE were given an IgE concentration value of 0. Comparison between the concentrations of IgE detected in the asthmatic samples compared to the control samples revealed no significant differences. This is likely due to the small number of samples with detectable IgE. However, samples Asthma_2 and Asthma_4 again had notably higher levels that the rest of the sample set. Within the asthmatic cohort it was these two subjects that had the highest levels of inflammatory proteins under investigation (see Supplementary Materials, S3), and thus the results of IgE quantification further support the concept of asthma sub-phenotypes with different circulatory immune status. As noted previously, such hypotheses require investigation with a much larger study cohort.

Overall, endotoxin levels were found to be reduced in the asthmatic subjects (**Figure 1E**; P value = 0.0650). Within the asthma cohort, subjects with additional atopic complications (i.e. allergic rhinitis, allergic dermatitis) displayed lower endotoxin concentrations compared to the asthmatic subjects that did not have additional atopic complications. This finding was further supported by the observation that within the control cohort, subjects with previously reported atopic dermatitis displayed circulatory endotoxin concentrations similar (i.e. lower than those subjects reporting no atopic conditions) to those observed in the asthma cohort.

### mRNA Sequencing and Differential Expression Analysis

Approximately 20,000,000 messenger RNA (mRNA) read pairs were generated from each plasma sample (average 44,000,000 ± 3,100,000 reads), with no significant differences in read count identified between the two cohorts.

Expression of a total of 14, 226 genes was detected through assessment of the circulating transcriptome (i.e. those RNAs present in the plasma). Given the nature of our sample type, the extent of read mapping to key mRNAs was confirmed visually by appraising the resulting BAM file against hg19 using IGV (data not shown). Sample Asthma_2 failed to map satisfactorily to hg19 and was thus excluded due to concerns this would induce bias into our downstream analyses. Statistical analysis, as detailed previously, revealed 287 genes were differentially expressed in the asthmatic subjects (as defined by a Q ≤ 0.05 and a Log2 Fold Change > 0.6). Within the asthmatic cohort, 90 of the differentially expressed genes showed significantly increased expression, and 197 genes displayed significantly decreased expression. Genes that displayed the highest degree of differential expression within the asthmatic subjects are listed in **Table 3**. A full list of differentially expressed genes can be viewed in the supplementary materials (Supplementary Materials, S4)

**Table 3:** Genes that were most differentially expressed in the asthmatic subjects (n = 4) compared to the control subjects (n = 5). Where genes are expressed in a condition-specific manner, Log2 fold change is replaced with “Control Only” or “Asthma Only” as appropriate. Quantity of the gene is shown as Fragments Per Kilobase of transcript per Million mapped (FPKM) reads

| Gene | Control Mean | Asthma Mean | Fold Change (log2) | Q Value |
| --- | --- | --- | --- | --- |
| **Downregulated Genes** |  |  |  |  |
| DOHH | 972.908 | 0 | <u>Control Only</u> | 0.002975 |
| PTRH2 | 87.7907 | 0 | <u>Control Only</u> | 0.002975 |
| C15orf41 | 79.1979 | 0 | <u>Control Only</u> | 0.002975 |
| HIST1H3I | 30.2331 | 0 | <u>Control Only</u> | 0.002975 |
| HOXC10 | 26.4924 | 0 | <u>Control Only</u> | 0.002975 |
| TSPYL5 | 18.9517 | 0 | <u>Control Only</u> | 0.002975 |
| NFXL1 | 17.8423 | 0 | <u>Control Only</u> | 0.002975 |
| RAB3IL1 | 15.1233 | 0 | <u>Control Only</u> | 0.002975 |
| LINC00085 | 15.0233 | 0 | <u>Control Only</u> | 0.002975 |
| ARV1 | 14.0641 | 0 | <u>Control Only</u> | 0.002975 |
| **Upregulated Genes** |  |  |  |  |
| HIST1H3C | 0 | 90.5782 | <u>Asthma Only</u> | 0.002975 |
| HDAC9 | 0.731644 | 52.1632 | 6.15575 | 0.005217 |
| PRAM1 | 0 | 3.05743 | <u>Asthma Only</u> | 0.005217 |
| PML | 0.948462 | 178.238 | 7.554 | 0.007164 |
| RAB6B | 0 | 8.90346 | <u>Asthma Only</u> | 0.007164 |
| NRP1 | 0.92425 | 18.8945 | 4.35354 | 0.010799 |
| CD93 | 0 | 14.3366 | <u>Asthma Only</u> | 0.010799 |
| GPR56 | 1.86976 | 98.5377 | 5.71975 | 0.012559 |
| MR1 | 1.07632 | 17.8916 | 4.0551 | 0.017952 |
| TOP1MT | 0.344555 | 59.0342 | 7.42067 | 0.017952 |

Interestingly, there were numerous genes that were expressed in a condition specific manner (96 genes were uniquely expressed in the control subjects, and 64 genes were uniquely expressed in the asthmatic subjects). To determine whether the asthmatic subjects had a distinct gene expression profile compared to the control subjects, genes that displayed robust levels of expression (as determined by a mean LOG2 FPKM score ≥ 6.0) were plotted as a heatmap and unsupervised cluster analysis was performed using Euclidean distance (**Figure 2**).

**Figure 2:**
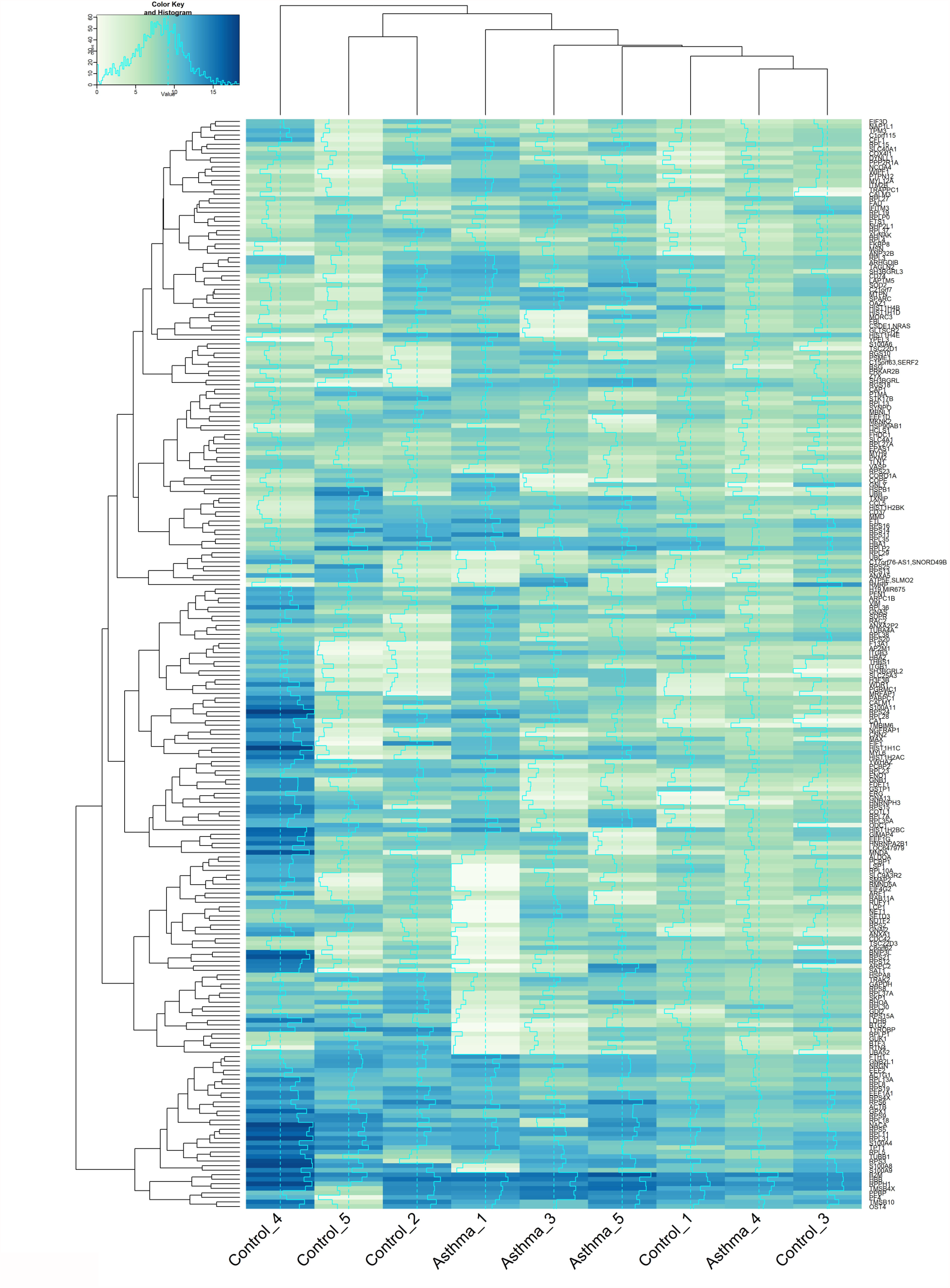
Heatmap showing highly expressed genes in control subjects (n = 5) and asthma subjects (n = 4). Gene expression is determined by quantification of circulatory mRNA present in the plasma samples and is expressed as log2 normalised Fragments Per Kilobase of transcript per Million mapped (FPKM) reads. Highly expressed genes, as determined by a mean log2 FPKM score ≥ 6.0 are plotted, and Cluster analysis (Euclidean distance) informs the X and Y-axis dendrograms.

Cluster analysis revealed that subject Control_4 had a relatively unique mRNA profile. For the remaining subjects, two clusters formed on the basis of circulatory mRNA populations. Cluster 1 was formed of Control_5 and Control_2; and Cluster 2 was comprised of Asthma_1, Asthma_3, Asthma_5, Control_1, Asthma_4, and Control_3. The dominance of asthmatic subjects in Cluster 2 suggests the possibility of a distinct asthma mRNA profile that would likely be more apparent in a larger sample group. Of interest, Asthma_4 clustered more closely with control Subjects Control_1 and Control_3. This asthmatic subject was the youngest member of the asthma cohort, with an age of 19 years, and the subject had been suffering from asthma for just 14 years compared to the mean length of 38 years that our other subjects had been living with the disease. It is tempting to speculate that asthmatic mRNA profiles become more divergent from control profiles as the disease progresses over time, however our sample size restricts further analysis of this.

The diversity of genes being expressed within the circulatory system was assessed using principal coordinate analysis (PCA) (**Figure 3**).

**Figure 3:**
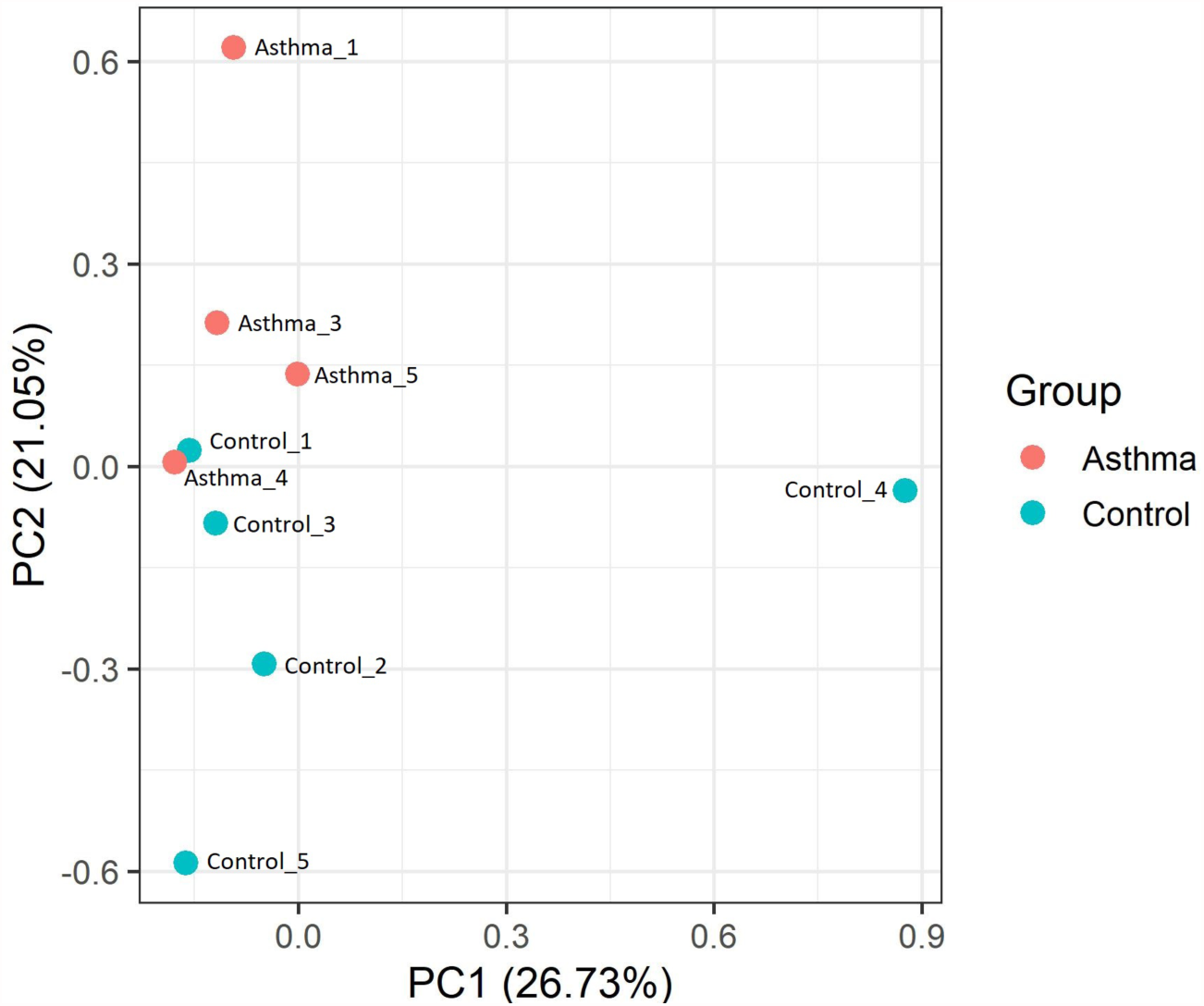
Principal component analysis ordination of Bray Curtis dissimilarity between circulatory mRNA populations present in control subjects (n = 5) and asthma subjects (n = 4). Principal component analysis was performed on a gene population dataset using quantitative mRNA Fragments Per Kilobase of transcript per Million mapped (FPKM) reads that had been normalised using log2. Only genes with mean FPKM scores ≥ 6.0 were included in the dataset, and the principal coordinate analysis was performed using Bray Curtis dissimilarity and R software. Blue data points = Control; Orange datasets = Asthma

Examination of Bray Curtis dissimilarity between the subjects found that, unsurprisingly, samples clustered similarly to that observed using unsupervised clustering (Euclidean distance). PCA analysis, did however reveal that the control and asthmatic subjects were differentiated on the basis of principal component (PC) 2, whereby asthmatic subjects had a positive PC2 score and control subjects had a negative PC2 score. Moreover, Asthma_4 clustered with the control subjects, thus providing additional evidence that this subject has a mRNA profile similar to the control subjects.

To determine whether differential gene expression could be linked to asthma pathology, we compared the differentially expressed genes identified herein, to a recently released database of genes associated with asthma pathology - AllerGAtlas, 2018 (41). Of the 287 genes identified as being significantly differentially expressed in the asthmatic subjects, 8 genes were identified in the asthma gene database. These genes included complement regulatory protein 46 (CD46), interleukin 7 receptor (IL7R), galactin 3(LGALS3), myeloperoxidase (MPO), neurotensin (NTS), phosphodiesterase 4A (PDE4A), toll-like receptor (TLR) 1, and vitamin D receptor (VDR). Four of the genes were upregulated in the asthmatic subjects (VDR, NTS, TLR1, and MPO) and four were downregulated in the asthmatic subjects (LGAL3, CD46, IL7R, and PDE4A) (**Table 4**). Moreover, gene expression was predominately condition specific. Of the upregulated genes, NTS, TLR1, and MPO mRNA was only detectable in the asthma samples, whilst in the downregulated genes, IL7R and PDE4R mRNA was only observed in the control samples (see Supplementary Materials, S4).

**Table 4:**
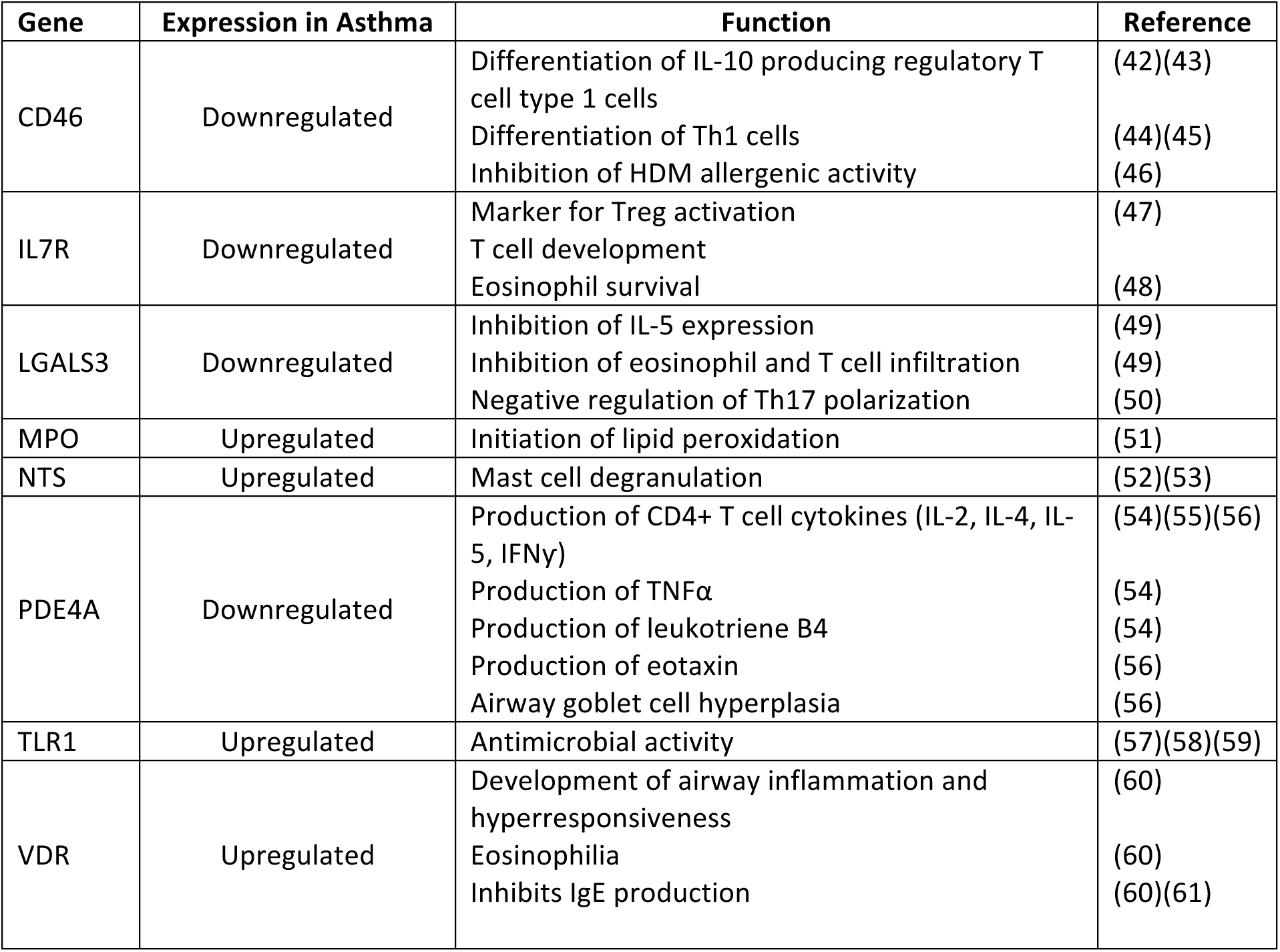
Genes with significant differential expression in the asthmatic subjects compared to control subjects that are associated with asthma pathology. Differential gene expression was determined using the Tuxedo protocol (Galaxy software) on log2 normalised mRNA Fragments Per Kilobase of transcript per Million mapped (FPKM) reads sequenced from plasma samples from asthma subjects (n = 4) and control subjects (n = 5). Gene function with regards to asthma pathology was determined using the asthma database AllerGAtlas, 2018 (41) and a general literature search using the relevant search engines.

The genes identified in the asthma gene database (41) were found to influence a number of key components of asthma pathology, including eosinophil and T cell migration, production of Th2 cytokines (IL-4, IL-5, and IL-13), mast cell degranulation, IgE production, and airway hyperresponsiveness. Moreover, several of the downregulated genes (CD46, IL7R), have been found to have roles in Treg differentiation and activation. These cells are important regulators of T cell activity (62–65), and thus downregulation of CD46 and IL7R suggests loss of control of T cell activity in the asthmatic subjects.

### miRNA Quantification

Approximately 10,000,000 micro RNA (miRNA) reads were generated from each plasma sample (range = 10,276,765 - 16,812,591, mean = 12,030,581 + 1,911,104), and there were no significant differences in read count identified between the control and asthma samples.

Using miRanalyzer (35) and edgeR (36), we identified 166 known miRNAs present in the plasma samples (**Figure 4**), which is consistent with previously reported studies (66–70). To determine whether the asthmatic subjects had distinct miRNA profiles compared to the control subjects, miRNA expression was plotted as a heatmap, and unsupervised clustering was performed using Euclidean distance (**Figure 4**).

**Figure 4.**
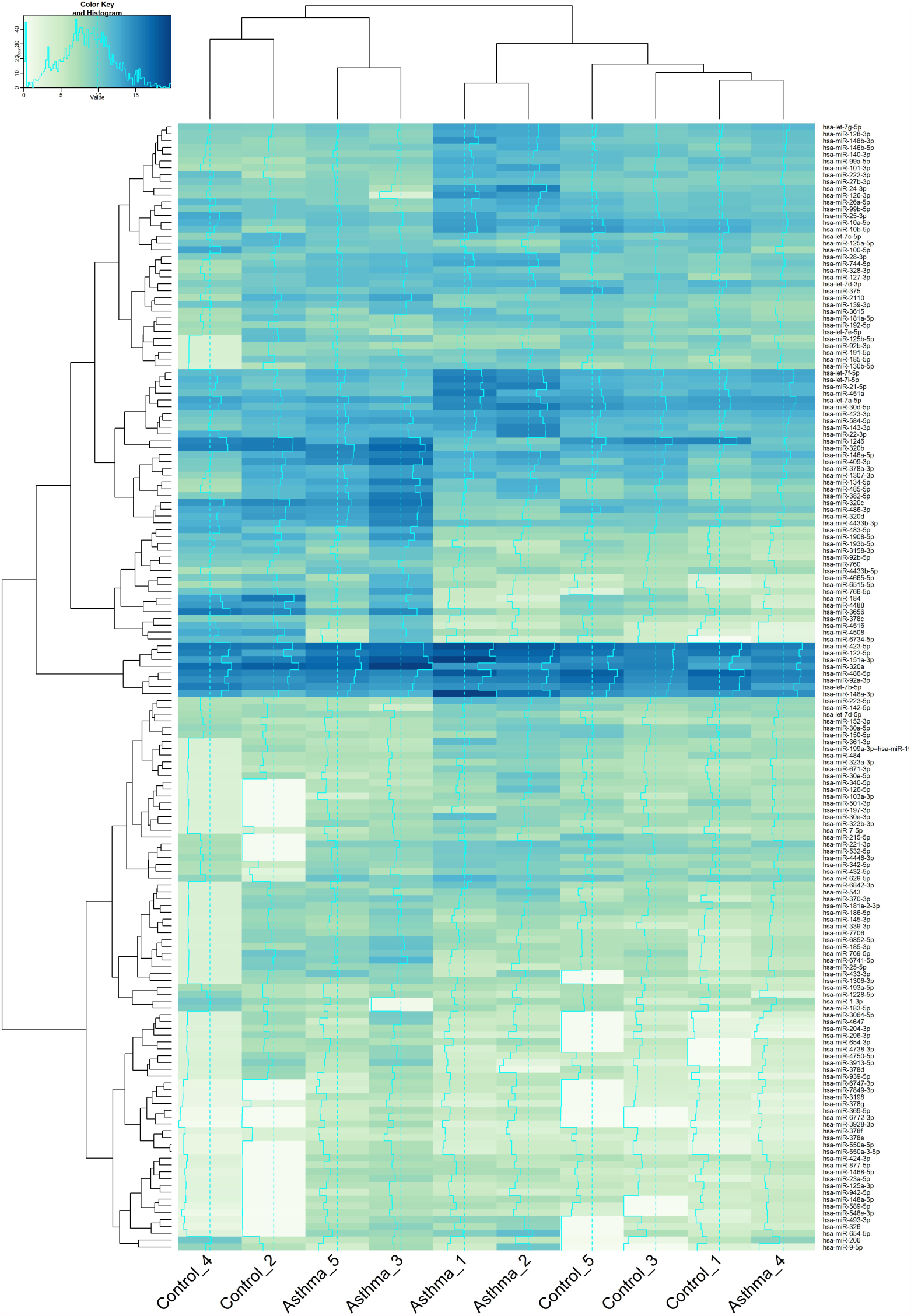
A Heatmap showing expression levels of circulatory miRNA in control subjects (n = 5) and asthmatic subjects (n = 5). miRNA expression is determined by quantification of circulatory miRNA detected in the plasma samples and is expressed as log2 normalised Counts per Million mapped (CPM) reads. Cluster analysis (Euclidean distance) informs the X and Y-axis dendrograms

Analysis of miRNA expression revealed the presence of two clusters with regards to the miRNA populations present within the plasma. Cluster 1 was composed Control_4, Control_2, Asthma_5, and Asthma_3; and Cluster 2 was made up of Asthma_1, Asthma_2, Control_5, Control_3, Control_1, and Asthma_4. Within each cluster two sub-clusters formed, and each sub-cluster was formed of either control subjects or asthma subjects. The one exception was Asthma_4, which clustered with other control subjects.

Of interest, the two asthma sub-clusters that formed appeared to be governed by the presence or absence of additional atopic complications. Asthma_5 and Asthma_3 clustered together and both subjects were free of additional atopic complication, whereas Asthma_1 and Asthma_2 clustered together, and both subjects had additional atopic complications such as allergic rhinitis. As we have noted previously, further study using a larger asthma cohort would be required to determine this association given the clear heterogeneity noted.

Statistical analysis revealed that 13 miRNAs were differentially expressed (defined as FDR P value ≤ 0.05 and a fold change ≥ 2.0) in the asthmatic subjects compared to the control subjects (**Figure 5**, see also Supplementary Materials S5). As predicted, Asthma_4 displayed miRNA levels similar to those observed in the control subjects. As stated previously, Asthma_4 was the youngest of the asthmatic subjects and had been living with the disease for the shortest period of time. It is tempting to speculate that asthmatic miRNA profiles become more divergent from control subjects as the disease progresses over time, and that this in turn alters gene expression.

**Figure 5:**
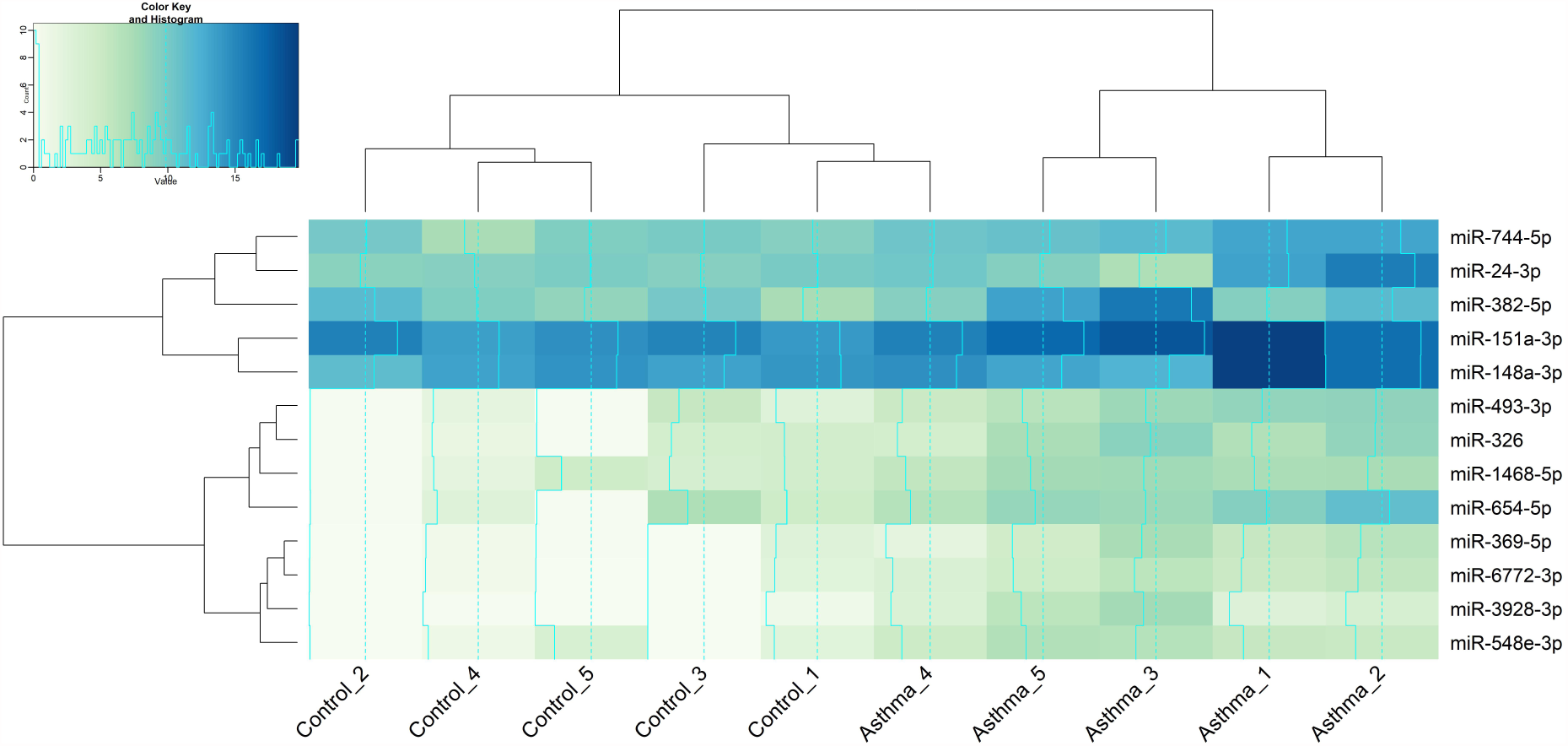
A heatmap showing expression levels of circulatory miRNA that displayed significant differential expression in asthmatic subjects (n = 5) compared to control subjects (n = 5). miRNA expression was determined by quantification of circulatory miRNA detected in the plasma samples and is expressed as log2 normalised Fragments Per Kilobase of transcript per Million mapped (FPKM) reads. Differential expression was determined using the edgeR program (Bioconductor software), and significant expression was defined as having a log fold change greater than 2.0 and a false rate of discovery (FDR) adjusted P value ≥ 0.05. Cluster analysis (Euclidean distance) informs the X and Y-axis dendrograms

### Functional Analysis

Causal inference analysis using Ingenuity Pathway analysis (IPA) software was performed to identify the likely upstream regulators responsible for the changes in mRNA and miRNA expression noted in the asthmatic subjects.

In total, 246 upstream gene regulators had a P value of overlap ≤ 0.05; indicating that they have altered functional activity in the asthmatic subjects on the basis of differential mRNA and miRNA expression. Of these regulators, seven had Z scores greater than 2.0, thus enabling their activity to be predicted. Two upstream regulators were predicted to have significantly increased activity in the asthmatic subjects (P value of overlap ≤ 0.05; Z score ≥ 2.0), and five were predicted to have significantly decreased activity asthmatic subjects (P value of overlap ≤ 0.05; Z score ≤ −2.0) in the (**Table 5**).

**Table 5:** Upstream gene regulators with predicted significantly altered activity in the asthmatic subjects (n = 4) compared to the control subjects (n = 5). Upstream regulators predicted to have significantly altered activity were defined as having a P value of overlap ≤ 0.05 and a Z score greater than 2.0. Activated upstream regulators are defined as having a Z score ≥ 2.0, and inhibited upstream regulators are defined as having a Z score ≤ −2.0. Target molecules activated = genes present in the RNA dataset that are activated by the upstream regulator; target molecules inhibited = genes present in the RNA dataset that are inhibited by the upstream regulator; target molecules affected = genes present in the RNA dataset whose activity is known to be altered by the upstream regulator but there is insufficient evidence to prove this is activation or inhibition.

| Upstream Regulator | Molecule type | Activity state | Z score | P value of overlap | # Target molecules activated | # Target molecules inhibited | # Target molecules affected |
| --- | --- | --- | --- | --- | --- | --- | --- |
| Sirolimus | Chemical drug | Activated | 2.75 | 0.0107 | 12 | 1 | 0 |
| GFI1 | Transcription regulator | Activated | 2.00 | 0.0077 | 4 | 0 | 1 |
| EIF4E | Transcription regulator | Inhibited | -2.00 | 0.0074 | 0 | 4 | 2 |
| Mycophenolic acid | Chemical drug | Inhibited | -2.00 | 0.0211 | 0 | 4 | 0 |
| Streptozocin | Chemical drug | Inhibited | -2.16 | 0.0492 | 0 | 5 | 1 |
| SOX4 | Transcription regulator | Inhibited | -2.24 | 0.0770 | 0 | 5 | 0 |
| SYVN1 | Transporter | Inhibited | -2.45 | 0.0069 | 0 | 6 | 0 |

Of interest, with regards to atopic asthma pathology, was the predicted activated state of GFI1, a transcription regulator induced by T cell activation and IL-4/STAT6 signalling. GFI1 is known to enhance Th2 expansion (71), and thus predicted activation of this transcription regulator would suggest increased T cell activation and subsequent expansion of the Th2 cell populations within the asthmatic cohort. This notion is further supported by the prediction of significant inhibition of the upstream regulator SOX4 in the asthmatic cohort. This transcription factor has been observed to suppress Th2 differentiation (72), and thus its inhibition would allow expansion of the Th2 populations within the asthmatic subjects. The predicted activated state of GFI1 would also influence innate immune responses within the asthmatic cohort. The transcription factor has been found to have a role in the development and maintenance of type 2 innate lymphoid cells (73); a cell population that has been found to be involved in allergic lung inflammation (74–76).

However, it should be noted causal inference analysis was performed on mRNA detected in the blood, and thus the cellular origins of the gene expression observed is unknown. Further study would be required to determine if GFI1 was indeed activated and SOX4 was inhibited in the relevant body sites and or relevant *in vitro* models of asthma pathology.

### Downstream Activity

Causal inference analysis using IPA was also used to predict the downstream consequences of the observed differential mRNA and miRNA expression within the asthmatic subjects. The downstream effects of the differential expression were primarily assessed by examination of the predicted canonical pathways and biofunctions impacted.

### Canonical pathway analysis

Fourteen canonical pathways were found to have significantly altered biological activity (P ≤ 0.05) within the asthmatic subjects (**Table 6**). In line with the findings of the upstream analysis, a number of canonical pathways involved in T cell and B cell activity, including signalling in rheumatoid arthritis, B cell development, and Nur77 signalling. It is interesting to note the canonical pathways involved in rheumatoid arthritis and Type 1 diabetes were identified, as both diseases have been found to display co-occurrence with asthma (77,78). It is tempting to speculate about the existence of similar / shared underlying immune pathologies in the three diseases.

**Table 6:**
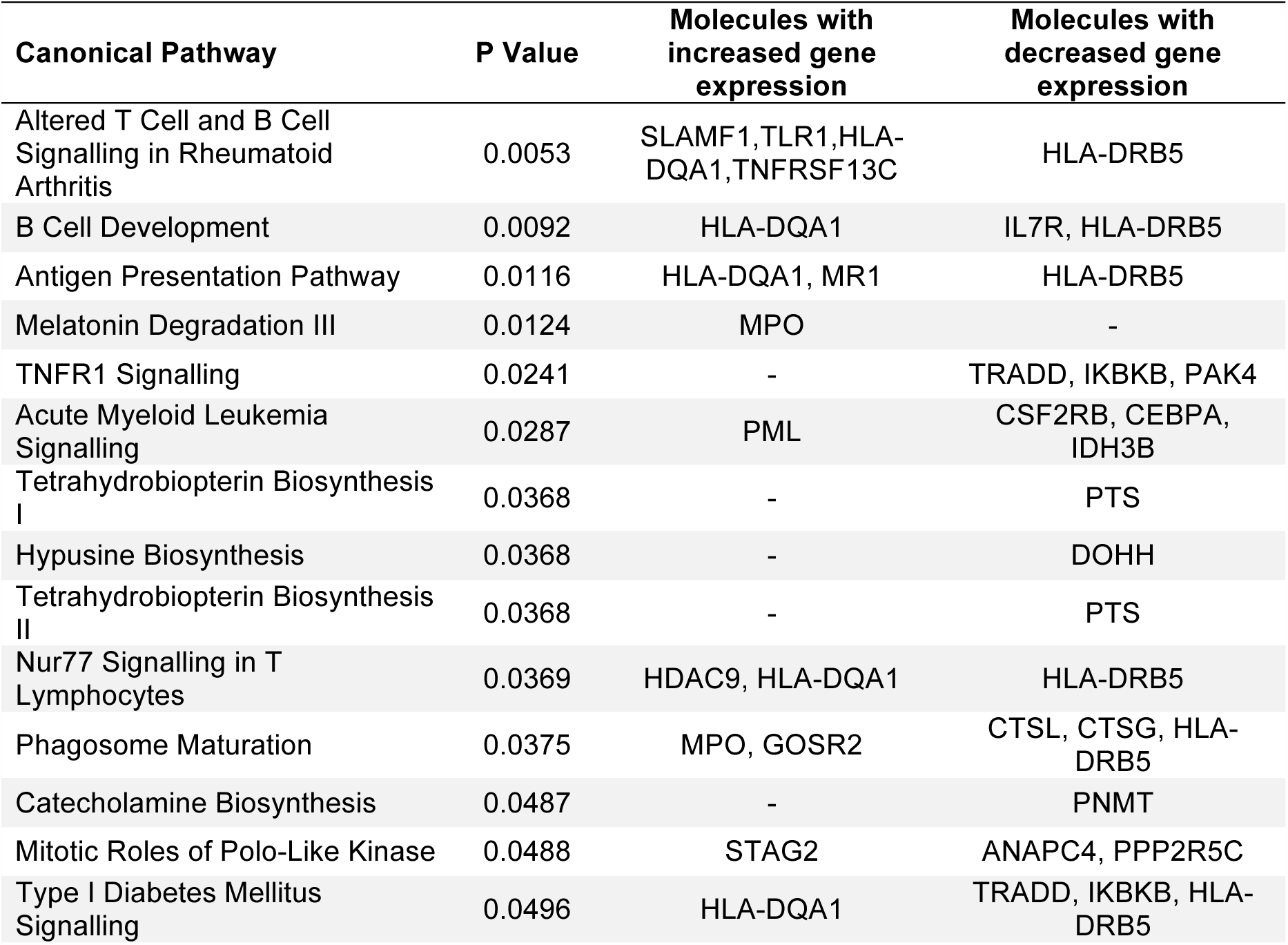
Canonical signalling pathways predicted to have significantly altered activity in the asthmatic subjects (n = 4) compared to the control subjects (n = 5). Casual interference using Ingenuity Pathway Analysis (IPA) software was used to predict downstream canonical signalling pathways likely to be affected by changes in gene expression and regulation in the asthmatic subjects. Molecules with increased gene expression are genes that had significantly increased numbers of mRNA reads in the asthma plasma samples, and molecules with decreased gene expression are genes that had significantly decreased numbers of mRNA reads in the asthma plasma samples. Canonical pathways that are defined as being significantly altered in the asthma subjects have a P value ≤ 0.05.

### Bio-function analysis

With regards to biological functions likely to be impacted by changes in the observed mRNA and miRNA expression patterns, a number of key immunological pathways were predicted to have altered activity within the asthmatic cohort (Table 7).

**Table 7:** Biological functions predicted to have significantly altered activity in the asthmatic subjects (n = 4) compared to the control subjects (n = 5). Casual inference using Ingenuity Pathway Analysis (IPA) software was used to predict biological functions likely to have altered activity in the asthmatic subjects. This was determined through analysis of genes and miRNA that had altered expression in the asthmatic subjects, to predict which biological functions would likely be altered. Biological functions predicted to be significantly altered in the asthmatic subjects were defined as having a P value ≤ 0.05 and a Z score greater than 2.0. Biological functions with predicted increased activity were defined as having a Z score ≥ 2.0, and biological functions with predicted decreased activity were defined as having a Z score ≤ −2.0

| Biological Functions | P Value | Activation State | Z score |
| --- | --- | --- | --- |
| Binding of endothelial cells | 9.75E-03 | Decreased | -2.123 |
| Binding of leukocytes | 1.73E-03 | Decreased | -2.062 |
| Cell transformation | 1.32E-03 | Decreased | -3.228 |
| Differentiation of fibroblast cell lines | 4.44E-03 | Decreased | -2.184 |
| Immune response of leukocytes | 6.79E-04 | Decreased | -2.031 |
| Interaction of endothelial cells | 3.55E-03 | Decreased | -2.346 |
| Killing of natural killer cells | 5.44E-03 | Decreased | -2.63 |
| Proliferation of hepatocytes | 6.53E-03 | Increased | 2.177 |
| Tumorigenesis of tissue | 4.94E-04 | Increased | 2.215 |
| Viral infection | 1.34E-02 | Decreased | -2.099 |

Altered activity was defined as having a P value ≤ 0.05 and a Z score ≥ 2.0 or ≤ −2.0; and in total 10 biological functions had significantly altered activity within the asthmatic subjects (**Table 7**).

Unsurprisingly, leukocyte activity was identified as being decreased in the asthmatic cohort. However, at this level of analysis, the downstream effects on biological function of the different classes of leukocytes was not determined, and thus further study would be required to ascertain which leukocytes would likely have altered activity in the asthmatic subjects as a consequence of the differential mRNA and miRNA expression. Study of the specific leukocyte classes affected by asthma would be crucial, as inhibition of the Th1 or Treg lymphocytes would likely enhance asthma pathophysiology, whereas inhibition of the Th2 lymphocytes would likely alleviate asthma pathophysiology.

It was also of interest to observe the predicted decrease in killing of natural killer cells. This cell population has been previously identified as having a critical role in immune defence against viruses and bacteria (79–82). In particular, viral infections have been long characterised to exacerbate asthma (83–86), and asthmatics have been observed to be deficient in type I IFN production (87–89), which likely influences natural killer cell activity. Moreover, in a murine model, natural killer cell activity was found to be decreased during a Th2 response (90). This suggests that in asthmatic subjects, as a consequence of a Th2 biased immune system, there is reduced natural killer cell activity, resulting in the known associations with asthma and respiratory infections. Moreover, this may also partially explain the changes in the airway microbiome we see in asthmatic populations.

### Characterisation of the Blood Microbiota

#### Bacterial Relative Abundance

Our previous characterisation (38) of the bacterial RNA present in the plasma samples found that the majority of bacterial RNA belonged to the Proteobacteria phylum (Total relative abundance = 83.9%; Control mean = 90.0%; Asthma mean = 80.3%), the Actinobacteria phylum (Total relative abundance = 7.5%, Control mean = 6.0%, Asthma mean = 7.5%), and the Firmicutes phylum (Total relative abundance = 6.6%, Control mean = 3.0%, Asthma mean = 9.0%) (**Figure 6**).

**Figure 6:**
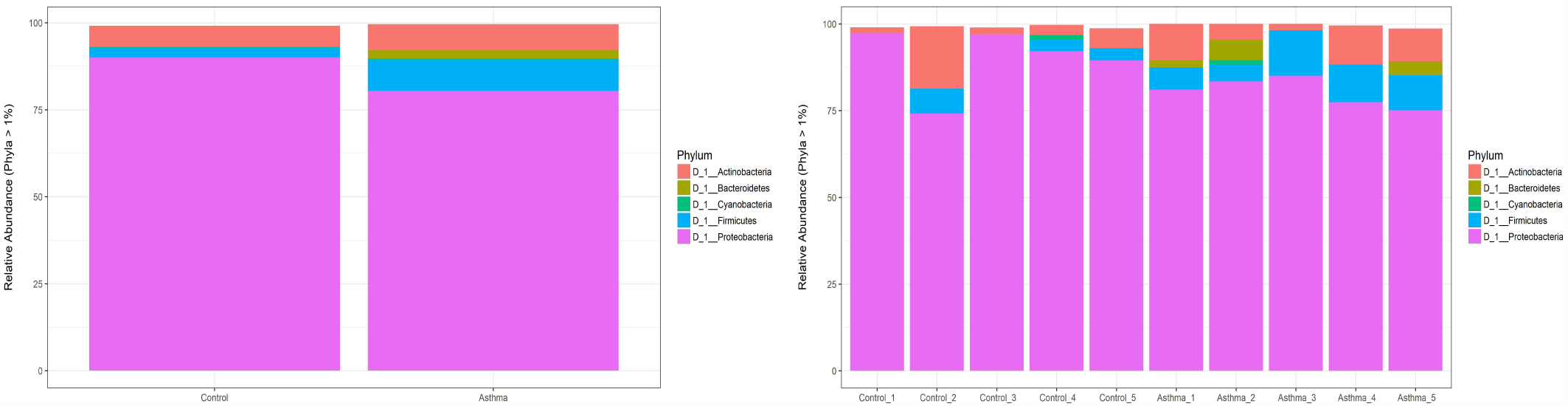
Microbial profile of the blood microbiome at the phylum level in asthmatic subjects (n = 5) and control subjects (n = 5). Composition of the blood microbiome was determined through sequencing of the bacterial V4 region of the 16S rRNA gene from bacterial DNA isolated from plasma samples from control subjects (n = 5) and asthmatic subjects (n = 5). The generated bacterial sequences were clustered (99% identity) in Operational Taxonomic Units (OTUs) to the Silva database and then assigned to bacterial taxonomic classes. **A** = microbial profile of the asthmatic subjects (n = 5) compared to the control subjects (n = 5). **B** = Microbial profiles of the individual plasma samples (n = 10)

In the asthmatic samples, 16S amplification and sequencing revealed a significant increase in Firmicutes (P value = 0.0148), associated with a concomitant decrease in Proteobacteria (P value = 0.0702) (**Figure 6**). To a lesser extent, members of the Bacteroidetes phylum were also detected in the blood samples, with increased levels of Bacteroidetes observed in the asthmatic subjects (Control mean relative abundance = 0.26%, range = 0.0 - 2.7%; Asthma mean relative abundance = 2.40%, range = 0 - 6.0%), although this was found to be non-significant increase (P value = 0.5258).

#### Lefse Analysis

Analysis of the phyla relative abundances detected in the blood was achieved using conventional statistical tests (unpaired *t* tests and Wilcox tests where appropriate) and suggested significant differences in the blood microbiome between control and asthma subjects. To test this, the linear discriminant analysis effect size (LefSe) method was applied to the 16S rRNA relative abundance data to determine the bacterial taxa most likely to explain the differences between the control and asthma blood microbiomes. LefSe was also used to determine the biological consistency and effect relevance of the observed differences in relative abundance.

In total, LefSe identified 8 bacterial taxa that showed statistically significant and biologically consistent differences in the asthmatic subjects compared to the control subjects (**Figure 7**). These findings were consistent with our previous analysis of the bacterial populations using standard statistical tests (data not shown). Six of the eight bacterial taxa displaying significant differences in relative abundance were increased in the asthmatic subjects, whilst 2 bacterial taxa were decreased. At the taxonomic class level, *Bacilli* were increased and *Bacteroidia* were decreased in the asthmatic subjects, whilst at the genus level both *Kocuria* and *Stenotrophomonas* were both increased in the asthmatic subjects.

**Figure 7:**
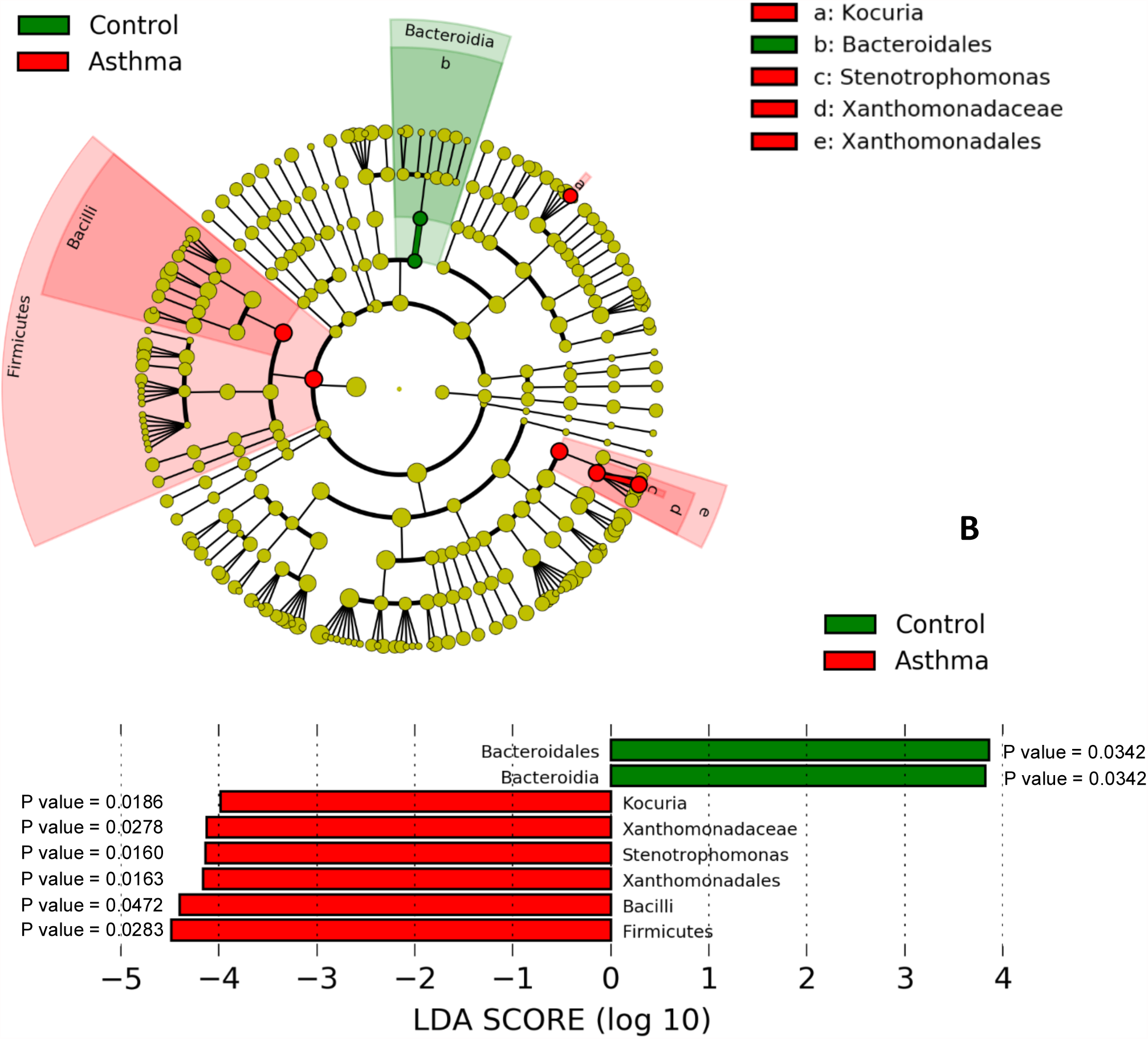
Comparison of the healthy blood microbiome (n = 5) and the asthmatic blood microbiome (n = 5) using LefSe. Linear discriminant analysis effect size (Lefse) analysis was performed on the bacterial taxa relative abundance values to determine the presence of bacterial taxa with statistically significant changes in abundance in the asthma blood microbiome compared to the control blood microbiome. **A.** Taxonomic cladogram showing control enriched taxa (Green) and asthma enriched taxa (Red). **B.** Effect size of the differential taxa. The control enriched taxa are indicated with a positive LDA score, and the asthma enriched taxa are indicated with a negative LDA score. The level of significance is indicated by the P value shown for each taxa.

The observed increases in Firmicutes were of particular interest as expansion of this phylum has been associated with severe asthma (21). Furthermore, increased levels of Firmicutes in the asthmatic subjects was predominately due to expansions of *Staphylococcus* and *Streptococcus* genera, both of which have been associated with the development of asthma during early childhood (91–94).

Additionally, our results were reflective of a previous study investigating the oral microbiome, whereby Firmicutes, *Stenotrophomonas*, and *Lactobacillus* were found to be increased in asthmatic subjects compared to the control subjects (95). This suggests that bacterial nucleic acid detected in the blood may have originated from the oral cavities, a theory that we consider in (38).

#### Bacterial Diversity

At the genus level, 81 bacterial genera were detected in the asthma plasma samples compared to 49 bacterial genera detected in the control plasma samples. Alpha and beta diversity of the bacterial populations present in the asthma and control groups was therefore assessed to determine whether there was significantly elevated bacterial diversity within the blood microbiome of the asthmatic subjects.

#### Alpha Diversity

Alpha diversity was determined by calculating the Chao1 index and Shannon index for each plasma sample. The control index scores were then compared to the asthma index scores to determine whether there were any significant differences between the two groups (Fig. 8).

**Figure 8:**
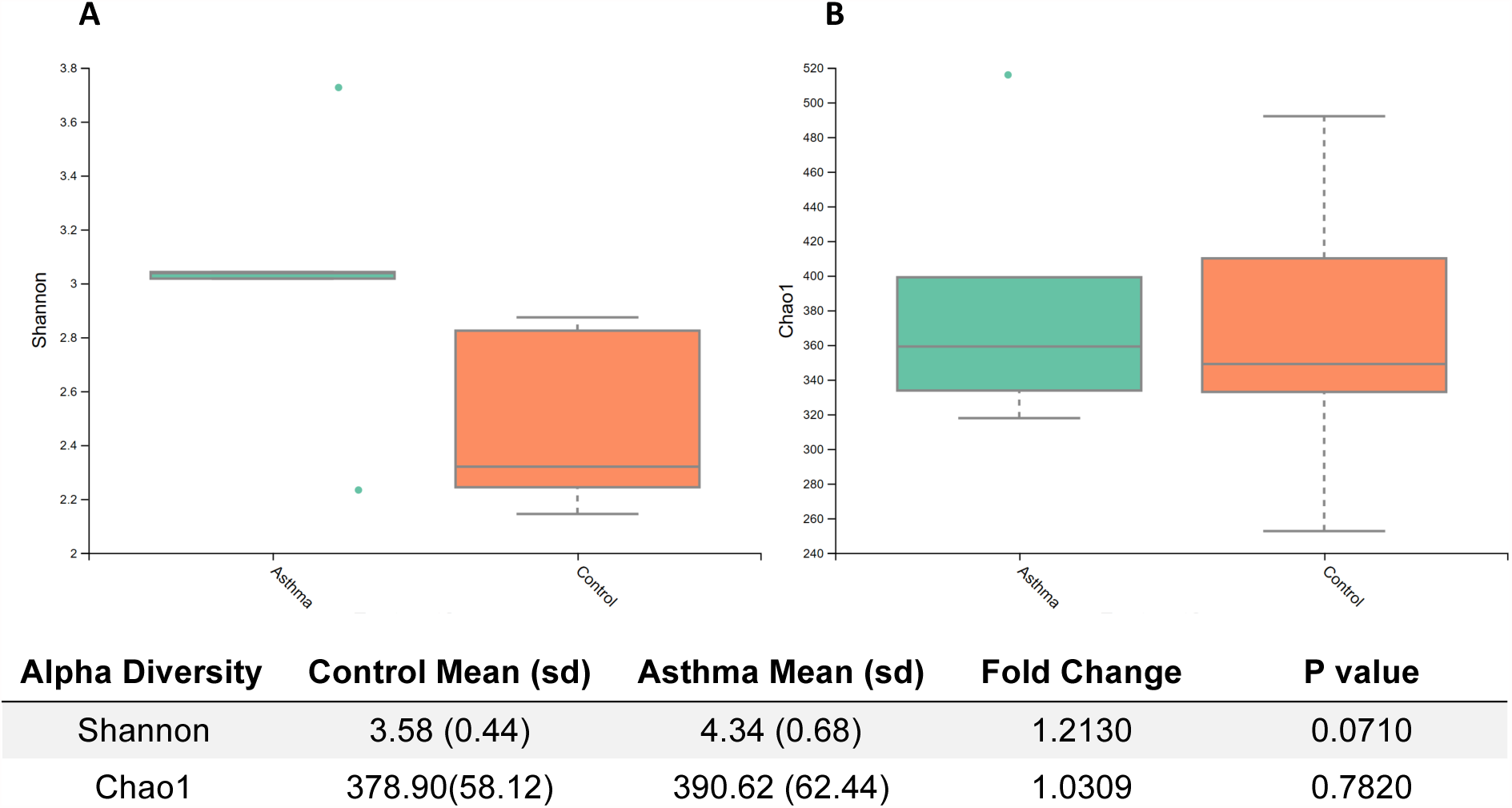
Comparison of alpha diversity present in the asthma blood microbiome compared to the control blood microbiome. Alpha diversity was measured using rarefied OTU tables generated from 16S rRNA sequencing data from plasma samples collected from asthma subjects (n = 5) and control subjects (n = 5). Shannon diversity index scores were generated from OTU tables in order to measure the richness of the plasma sample and evenness of bacterial taxa present in the sample. Chao1 index scores were measured to determine the predicted number of bacterial taxa present in the plasma samples by extrapolating out the number of rare organisms that may not have been detected due to under-sampling. **A** = Comparison of Shannon index scores generated from asthma plasma samples (n = 5) and control plasma samples (n = 5), **B =** Chao1 index scores generated from asthma plasma samples (n = 5) and control plasma samples (n = 5).

Comparison between the asthma and control cohorts revealed that the asthmatic subjects scored higher Chao1 and Shannon index scores than the control subjects, thus suggesting that asthma is associated with increased bacterial diversity (**Figure 8**). This was particularly apparent for the Shannon diversity scores (P value = 0. 0710) (**Figure 8**). Intriguingly, one of the asthma subjects, Asthma_3, displayed a Shannon diversity score more similar to the controls than the other asthmatic subjects. This subject developed asthma relatively late in childhood (age 12 years), and so it is possible that the age of asthma onset may influence the level of microbial diversity present in the blood. This is further supported by the high levels of alpha diversity present in the blood of Asthma_5, an asthmatic subject who was diagnosed with asthma early on in childhood (3 years).

#### Beta Diversity

Beta diversity was calculated to determine how similar the blood samples were to one another with regards to bacterial diversity. This enabled not only comparison between the asthma and control subjects, but also between the different members within each group.

Beta diversity was determined by performing principal coordinate component (PCoA) analysis using weighted UniFrac distances (**Figure 9**). PCoA analysis found that beta diversity was principally a consequence of PCo1 variation (37.1%), and overall the asthmatic subjects had higher PCo1 values with regards to beta diversity within the blood microbiome compared to the control subjects.

**Figure 9:**
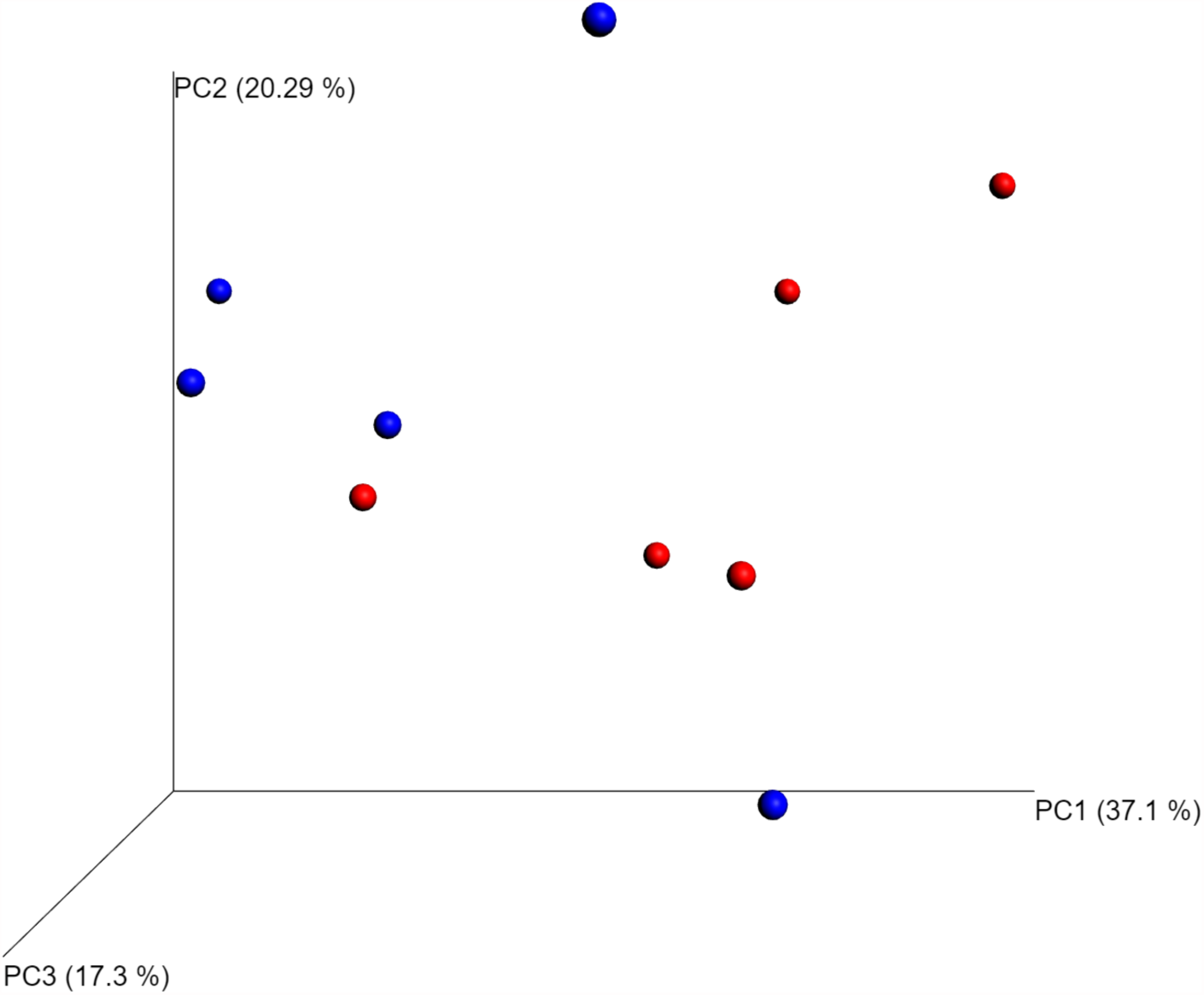
Beta diversity of the blood microbiome from asthmatic subjects (n = 5) and control subjects (n = 5) using weighted UniFrac distance. Principal coordinate analysis (PCoA) was performed on OTU tables generated from 16S rRNA sequencing data from plasma samples collected from asthma subjects (n = 5) and control subjects (n = 5). Quantitative phylogenetic distances between each of the samples was measured using a weighed UniFrac distance matrix, and the weighted UniFrac distances were plotted as a PCoA graph to show beta diversity within plasma samples from control subjects (n = 5; data plots = blue) and asthma subjects (n = 5; data plots = red)

#### Concluding Remarks

This study aimed to characterise a small yet specific population of HDM-sensitive adult asthma patients who had developed asthma during childhood. A range of molecular techniques was applied to characterise gene expression and regulation, inflammatory protein levels, and nucleic acid evidence of bacteria present in the blood. This was carried out in an effort to increase our understanding of this particular asthma phenotype, to begin to explore the molecular mechanisms responsible, and to identify any candidate biomarkers for further study.

At the protein level, the asthmatic subjects displayed increased inflammatory protein levels in the blood compared to the control subjects. This was particularly apparent for GM-CSF, IFN and TARC. The range of inflammatory protein levels within the asthmatic subjects was noticeably higher than the range observed for the control subjects. This was explained by the presence of two distinct clusters in the asthmatic cohort; cluster one was composed of subjects Asthma_2 and Asthma_4, and was characterised by high inflammatory protein levels; and cluster two, composed of Asthma_1, Asthma_3, and Asthma_5, and characterised by lower levels of inflammatory proteins. An association between the existence of other atopic complications, in particular evidence of atopic dermatitis, and IL-17A levels was unexpectedly observed.

Measurement of total IgE concentration within the blood revealed that IgE was detectable in half of the subjects under investigation (3 control subjects and 2 asthmatic subjects) and was significantly increased in the asthmatic subjects, when detected. The low detection rate of IgE was not unexpected given its short half-life (approximately two days) and low concentration levels within the blood (96). IgE was detected in asthma subjects belonging to the proposed cluster one, and this further supports the theory of asthmatic subjects forming sub-phenotypes on the basis of circulatory inflammation. In contrast to IgE, endotoxin levels were decreased in the asthmatic subjects (P value = 0.0650), and there appeared to be an inverse correlation between circulatory endotoxin levels and the reporting of additional atopic complications. This was a particularly interesting finding as exposure to endotoxin during early childhood has been previously found to be protective of the development of childhood asthma (97–100), and we were able to detect changes in endotoxin levels in our adult cohort.

Analysis of the diversity of RNA expression within the blood revealed that our asthmatic donors had more similar RNA profiles to one another than they did to the control subjects; this was particularly apparent in the miRNA analysis. When combined with our differential expression analyses, we identified specific mRNA and miRNA populations within the blood that were distinct between the healthy and disease states. Interestingly, asthma severity and the use of anti-inflammatory medication appeared to further influence RNA profiles although we note the limitations of our sample size, and acknowledge the need for a larger sample size to explore this phenomenon fully. With regards to the unmet need for asthma biomarkers, we identified various mRNAs in the circulation that were expressed in a condition-specific manner, including HIST1H3C, PRAM1, RAB6B and CD93. Of these, elevated levels of soluble CD93 have been previously reported in the serum of asthmatics during acute asthma exacerbations (101) and in the serum of steroid-naïve asthmatic patients (102).

Our microbial characterisation informed by 16S rRNA amplification and sequencing, revealed increased levels of Firmicutes and decreased levels of Proteobacteria within the blood of our asthmatic donors. This finding was accompanied by increased bacterial diversity within the blood of asthmatic subjects, and the identification of several additional bacterial taxa displaying significantly altered levels dependent on disease state. The observed decrease in circulating Proteobacteria rRNA in the asthmatic state is thought to be indicative of reduced Proteobacteria carriage within the asthmatic subjects at a distant microbiome niche (e.g. the gut, airways and oral cavity). This may explain the decreased levels of endotoxin (protein) detected in our asthmatic subjects, given that endotoxin-producing gram-negative bacteria dominate this phylum. Previous studies have associated childhood asthma and reduced endotoxin exposure, and it is interesting to note that we detected this same phenomenon in our adult asthma cohort, many years following childhood. Furthermore, our asthma patients were found to have increased levels of Bacteroidetes rRNA, and this appeared to be dependent on medication status with those patients taking anti-inflammatory medications having lower levels of circulating Bacteroidetes 16S rRNA than those who were not. As blood circulates the body and functions as a medium that samples from virtually all body sites (103), it was not possible to determine herein the microbial niche from which these signals originated. That said, we hypothesise that changes in the blood are reflective of dysbiosis at distant site(s) with well-characterised microbial communities (e.g. the gut, oral cavity and skin), and have significant biomarker potential.

This study provides a valuable insight into the systemic changes evident in the HDM-associated asthma, identifies a range of molecules that are present in the circulation in a condition-specific manner (with clear biomarker potential), and highlights a range of hypotheses for further study. Moreover, our data also provide an insight into the level of heterogeneity observed both within the control and asthma samples investigated, and will be of use for informing sample size calculations for future studies.

## Research Affiliations

The National Institute for Health Research Health Protection Research Unit (NIHR HPRU) in Health Impact of Environmental Hazards at King’s College London in partnership with Public Health England (PHE) in collaboration with Imperial College London

## Acknowledgements

This research was part funded by the National Institute for Health Research Health Protection Research Unit (NIHR HPRU) in Health Impact of Environmental Hazards at King’s College London in partnership with Public Health England. The views expressed are those of the author(s) and not necessarily those of the NHS, the NIHR, the Department of Health or Public Health England.

## Author Contributions

DPT, MOL and TWG conceived the original study. DPT developed and refined the molecular approach used. EW conducted the laboratory work. EW and DPT conducted the data analysis. EW and DPT interpreted the original data. EW and DPT prepared the original manuscript. EW, MOL, TWG and DPT reviewed and approved the manuscript.

